# A theoretical model of neural maturation in the developing spinal cord

**DOI:** 10.1101/668640

**Authors:** Piyush Joshi, Isaac Skromne

**Author notes:** **Corresponding author:** (I.S.).

## Abstract

Cellular differentiation is a tightly regulated process under the control of intricate signaling and transcription factors interaction network working in coordination. These interactions make the systems dynamic, robust and stable but also difficult to dissect. In the spinal cord, recent work has shown that a network of FGF, WNT and Retinoic Acid (RA) signaling factors regulate neural maturation by directing the activity of a transcription factor network that contains CDX at its core. Here we have used partial and ordinary (Hill) differential equation based models to understand the spatiotemporal dynamics of the FGF/WNT/RA and the CDX/transcription factor networks, alone and in combination. We show that in both networks, the strength of interaction among network partners impacts the dynamics, behavior and output of the system. In the signaling network, interaction strength determine the position and size of discrete regions of cell differentiation and small changes in the strength of the interactions among networking partners can result in a signal overriding, balancing or oscillating with another signal. We also show that the spatiotemporal information generated by the signaling network can be conveyed to the CDX/transcription network to produces a transition zone that separates regions of high cell potency from regions of cell differentiation, in agreement with most *in vivo* observations. Importantly, both networks have built in robustness to extrinsic disturbances. This analysis provides a model for the interaction conditions underlying spinal cord cell maturation during embryonic axial elongation.

## Introduction

Cells sequentially differentiate from high to low potency states, under the guidance of extracellular signals working in coordination with intracellular transcription factors. Signals regulate the individual and network activity of the transcription factors by providing spatial and temporal information [1–4]. In turn, transcriptional network dictates a cell’s competence and response to extracellular signals [5–7]. Because signaling information changes the composition of a cell’s transcriptional components, this creates an intricate and dynamic cross-regulatory system for guiding cell differentiation that has been challenging to untangle and comprehend [1, 3, 4].

Vertebrate spinal cord provides an advantageous model to study the cross-regulatory dynamics involved in central nervous system development in particular, and differentiation in general. The head (rostral) to tail (caudal) development of spinal cord during vertebrate body extension results into a characteristic spatial separation of temporal differentiation events [8–10], facilitating the study of their regulation. Briefly, the spinal cord neural progenitors (NPs) are derived from a bipotent population of cells located at the caudal most end of the embryo, the neuro-mesodermal progenitors (NMPs). In the early embryo, the region where NMPs reside is known as the caudal lateral epiblast and node streak border, and in the late embryo, the caudal neural hinge [8–10]. During development, NP cells exit the NMP domain rostrally and then, sequentially, transit through different maturation states as they become part of the spinal cord [8, 11, 12].

NP cell maturation is driven by synergistic and antagonistic interactions between the signaling factors FGF, WNT and Retinoic Acid (RA), turning on and off key transcription factors required for caudal-to-rostral maturation events (**Fig 1A**). Current models propose that two opposite signaling gradients regulate spinal cord cell maturation [13]: from caudal/high to rostral/low, FGF and WNT gradients prevents cell differentiation by promoting high potency cell states caudally; whereas an opposite rostral/high to caudal/low gradient of RA secreted from somites promotes cell differentiation rostrally. Importantly, FGF and WNT activity gradient counteract RA activity gradient. In this way, cells located caudally experience high levels of FGF/WNT and no RA, which drives expression of bipotency markers *T/Bra*, *Sox2*, and *Nkx1.2* [11, 14, 15]. T/BRA and SOX2 are transcription factors that repress each other and promote different cell fates, with T/BRA promoting mesoderm and SOX2 promoting neural fates [16–19]. In addition, both transcription factors downregulate FGF and WNT, initiating the early differentiation of mesoderm or neural tissues [16]. NMPs that continue to transcribe *Sox2* but not *T/Bra* assume NPs identity and become part of the growing neural plate. As NPs transit through the maturing neural plate, they experience a further gradual loss in FGF and WNT, and a gradual increase in RA signaling. This new environment lead to the caudal-to-rostral downregulation of a third bipotency marker, *Nkx1.2*, and upregulation of the early differentiation gene *Pax6* [20]. Subsequently, under RA regulation, PAX6 activates late differentiation genes such as *Ngn2* (**Fig 1B**) [10, 21, 22]. Recently, we experimentally mapped the interactions of these transcription factors into a gene regulatory network (GRN) and place it in the context of the FGF/WNT-RA signaling network (**Fig 1C**) [12]. This work identified the transcription factor CDX4 as a core system component essential for the sequential maturation of NPs into mature neuronal precursors (**Fig 1C**).

**Fig 1.**
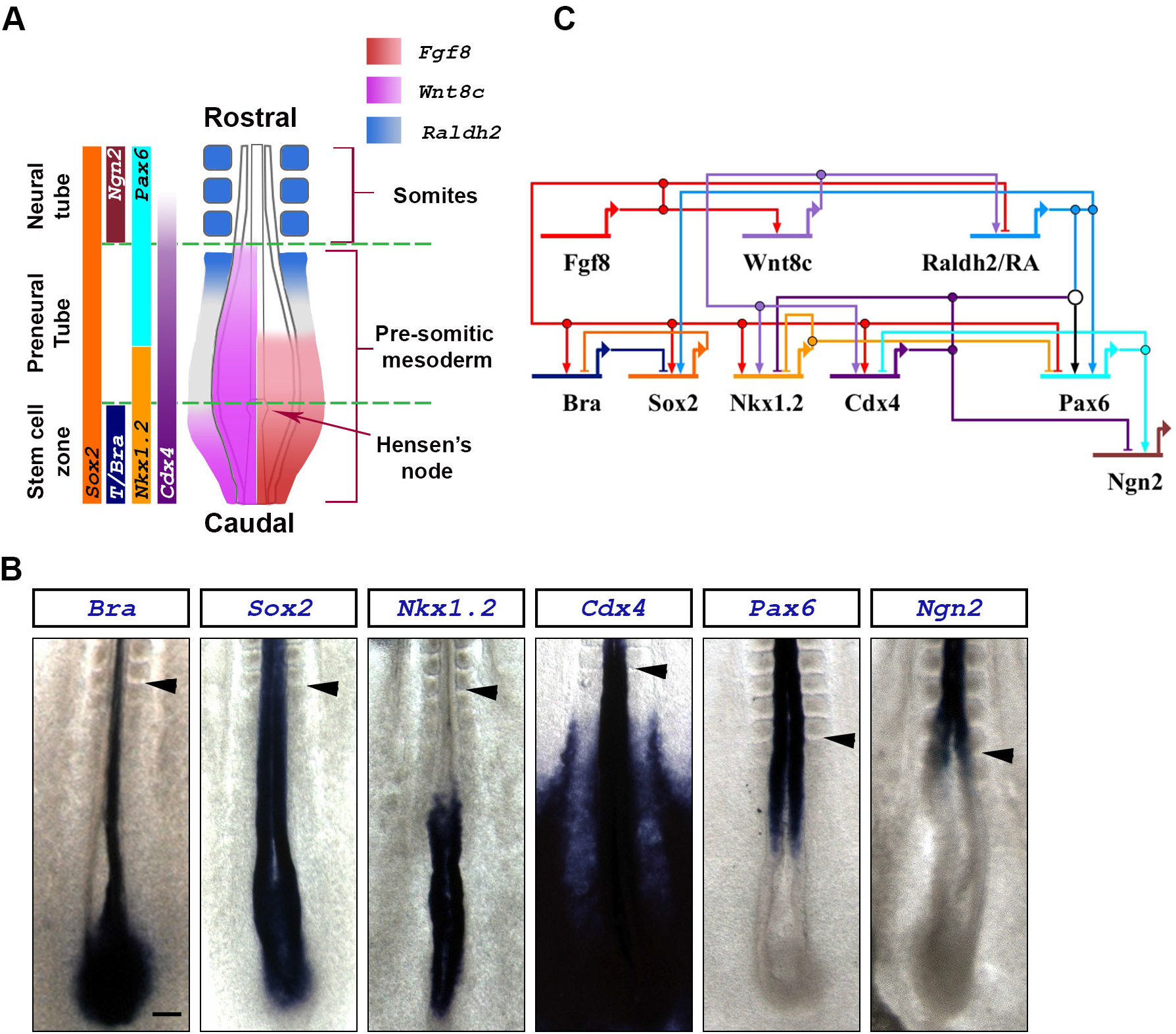
Expression domains and network interactions of key signaling and transcription factors involved in caudal spinal cord maturation. (**A**) Schematic representation of the caudal end of a stage HH10-11 chick embryo (dorsal view). Expression domains of *Fgf8* (red) and *Wnt8c* (magenta) signaling factors, and the Retinoic Acid synthesizing enzyme *Raldh2* (blue), are superimposed on the diagram (based on [23]). Expression domain of relevant transcription factors are indicated on the left (based on [12]). (**B**) Expression domains of key transcription factors involved in caudal spinal cord maturation. Embryos are stage HH10-11. Scale bar is 200μm. Arrowheads indicates the anterior boundary of the last formed somite. Transcription of the *T/Bra* gene along the embryo’s midline is in the notochord underlying the neural tissue, where it is absent. (**C**) Postulated gene regulatory network showing interaction between signaling and transcription factors (based on [12, 16, 21, 23]).

Here we use partial and ordinary (Hill) differential equations to dynamically analyze the GRN driving NP cell maturation during early spinal cord development. As the transcription factor network depends upon inputs form the FGF-WNT-RA signaling network, we first analyzed the postulated effectiveness of the signaling network to work as a signaling switch [23]. We then used the resulting signaling dynamics as input to evaluate the performance of the underlying transcription GRN in its ability to generate cell state patterns similar to those observed in experimental models. Our results show that signaling interaction can give rise to various developmentally observed phenotypes based on a limited subset of interaction parameters, and these behaviors are robust and stable to perturbations. These robust behaviors are due to strong cross-regulation interactions between system elements. Our results suggests that the dominant predictor of the GRN response is the interaction strength among network partners. By outlining the conditions that permit the operation of the GRN during NP maturation *in silico*, the model predicts and informs on cellular behaviors of the system *in vivo*.

## Methods

Hill equations were used to model the signaling and transcription factor interaction networks. Within each equation, Hill constants were used to vary the strength of interaction between a molecule and its target substrate (e. g., interaction between a transcriptional regulator and its target gene; **Fig S1**). We first modeled the signaling network as it is responsible for the spatial regulation of the transcription factor network. Partial differential equation were used to model the synthesis, diffusion, degradation and interactions between FGF, WNT and RA signals. The output of the signaling network was then used as the input for the transcription factor network. This transcription factor network was solved using ordinary differential equations. We used MATLAB (MathWorks, Natick, MA) to solve the equations numerically and to plot the simulations. Partial differential equations were solved using *pdepe* solver and ordinary differential equations using *ode45* solver.

### Hill equation based interaction model

In both networks, the rate of mRNA change was modeled using ordinary differential equations, and the rate of protein change was modeled using either ordinary differential equations for transcription factor or partial differential equations for signaling factors to accommodate diffusion [24–26]. These equations follow the general form;

mRNA dynamics:

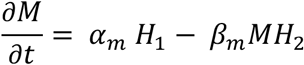

Protein dynamics for transcription factors:

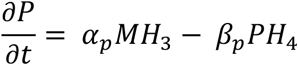

Protein dynamics for signaling factors:

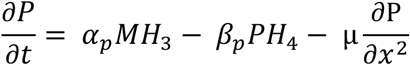

,where,

*α*_*m*_ = Transcription rate constant. *α*_*p*_= Translation rate constant.

*β*_*m*_= mRNA decay rate constant. *β*_*p*_= protein decay rate constant.

*M* = mRNA concentration. *P* = protein concentration.

μ = diffusivity coefficient. *x* = spatial dimension.

*H*_1_, *H*_2_, *H*_3_, *H*_4_ are independent Hill functions that can describe one of four general types of regulatory interaction between network components (see Supp. Methods): (1) inductive interactions from one or multiple activators, (2) repressive interactions from one or multiple repressors, (3) coordinated interactions between activators and repressors binding to separate regulatory sites and (4) competitive interactions between activators or repressors binding to the same regulatory sites. For each factor being modeled, we replaced *H*_1_, *H*_2_, *H*_3_, and *H*_4_ functions with the appropriate equation that best describes the regulatory interactions observed experimentally.

### Equations modeling the signaling interactions network

Partial differential Hill equations were used to model FGF8, WNT8C and RA network of interactions (**Fig 2A**). These interactions were modeled within a spatial maturation domain restricted to a 2500 microns extending from the NMP zone to the anterior boundary of the last formed somite (stage HH10-11 embryos; **Fig 1B, 2B**). This spatial maturation domain moves caudally and in synchrony with the NMP zone during axial elongation, thus appearing stationary with respect to the NMP zone (**Fig 2B**). When available, we used parameter values that have been determined experimentally, within reported ranges. For parameters that have not been determined experimentally (e.g., rates constants for mRNA and protein synthesis and degradation), we used parameters values comparable to those used in other models [27–29].

**Fig 2.**
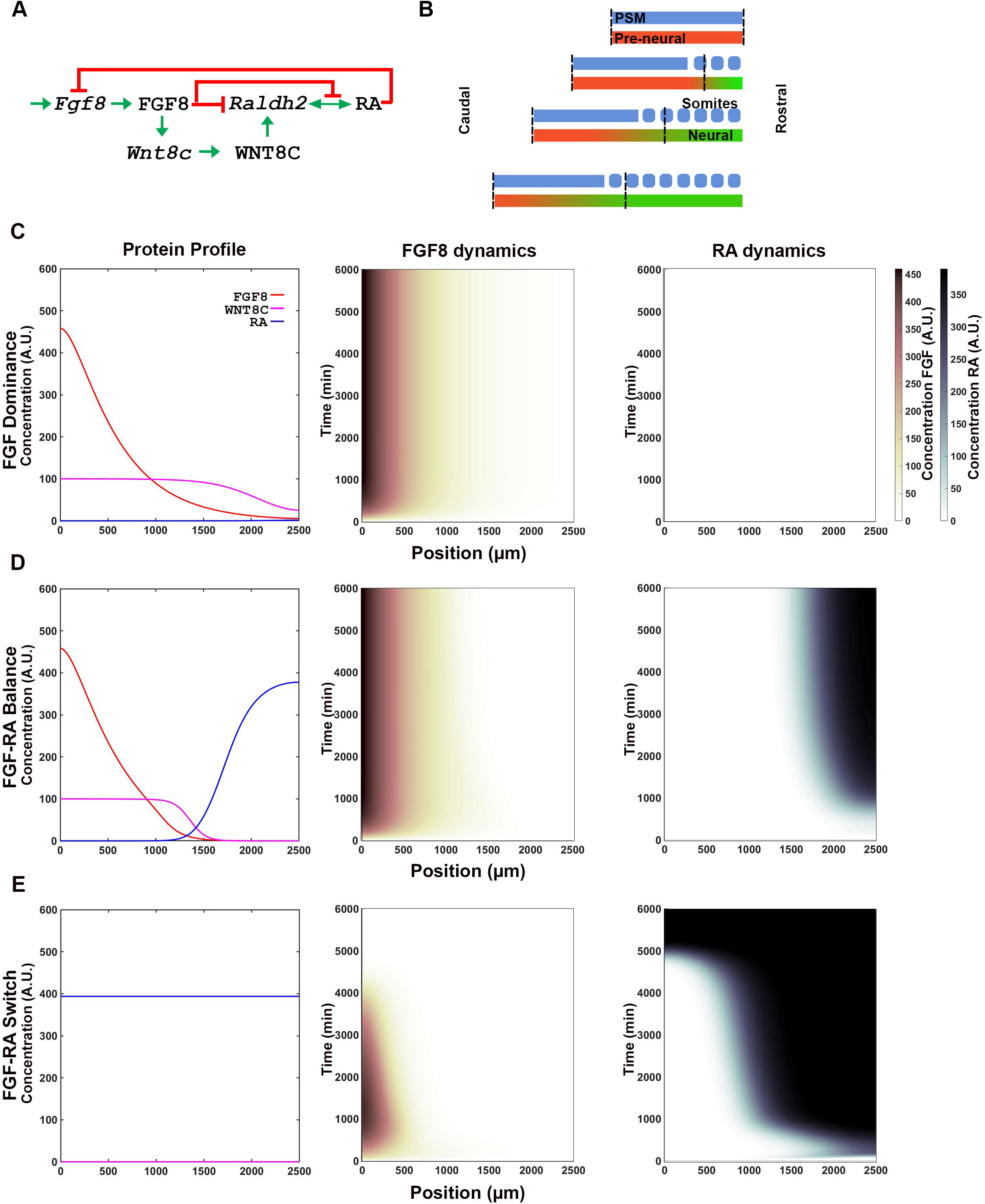
Signaling network output is determined by the strength of interactions between FGF, WNT, and RA pathway components. (**A**) FGF8, WNT8C and RA signaling pathway interaction network based on [23]. Names in lower case indicate mRNA and upper case proteins (FGF8 and WNT8C) or metabolites (RA). (**B**) The spatial maturation domain where the signaling network operates extends from the NMP cells to the anterior boundary of the last formed somite (vertical dashed lines; x-axes on graphs). The domain in the simulation has a constant length maintained by a caudal movement that is equivalent to the rate of NMP cell proliferation. Initially undifferentiated cells differentiate at a rate defined by the simulation (red to green transition). (**C-E**) Representative FGF8 dominant (C), FGF-RA balance (D), and FGF-RA switch (E) simulation profiles obtained using parameters shown in **Table 1**. Left graphs shows the levels of the signaling molecules FGF8 (red), WNT8C (magenta) and RA (blue) across the maturation domain at the end of the simulation (t=6000 min; arbitrary units AU). Center and right graphs show heat maps of FGF8 (center) and RA (right) accumulation in the maturation domain (x-axis) over time (y-axis). AU scale for FGF (maroon gradient) and RA (blue gradient) are shown at right of graphs.

**Table 1.** Examples of Hill constants combinations tested to investigate signaling dynamics behavior.

|  |  |  |  |  |  |  |  |  |
| --- | --- | --- | --- | --- | --- | --- | --- | --- |
| Hill constants | I | II | III | IV | V | VI | VII | VIII |
| $F_{FW}$<br>FGF dependent activation<br>of *Wnt8c* transcription | 10 | 10 | 10 | 10 | 10 | 10 | 10 | 10 |
| $F_{FR1}$<br>FGF dependent repression<br>of *Raldh2* transcription | 1 | 1 | 5 | 10 | 10 | 10 | 2 | 20 |
| $F_{FR2}$<br>FGF dependent activation<br>of RA degradation (via<br>CYP26A enzymes) | 2 | 10 | 15 | 10 | 10 | 10 | 20 | 20 |
| $W_{WR}$<br>WNT dependent<br>activation of *Raldh2*<br>transcription | 1 | 1 | 0.2 | 0.5 | 0.2 | 0.5 | 1 | 1 |
| $R_{RF}$<br>RA dependent repression<br>of *Fgf8* transcription | 10 | 1 | 0.2 | 0.1 | 0.3 | 0.45 | 1 | 20 |
| $R_{RR}$<br>RA dependent activation<br>of *Raldh2* transcription | 50 | 50 | 50 | 50 | 50 | 50 | 300 | 300 |
| Outcome (t=6000 min) | FGF<br>dominant | FGF-RA<br>balance | FGF-RA<br>switch |  |  |  | RA aberrant/<br>oscillatory |  |

#### FGF8 PRODUCTION

*Fgf8* transcription is restricted to the NMP zone through positive autoregulatory loops and inhibitory signals [30]. FGF8 indirectly stimulates its own transcription by inducing transcription of *Nkx1.2*, *Cdx*, and WNT/ß-catenin pathway components [30]. RA secreted from somites restricts *Fgf8* to the NMP zone in a concentration-dependent manner [13]. RA inhibition is excluded from NMP zone by CYP26A, an RA-catabolizing enzyme whose gene is activated by FGF8 [31]. We simulated *Fgf8* positive autoregulatory loop by assuming a basal exponential level of gene transcription, and its restriction to the caudal end of the spatial maturation domain by allowing RA to decrease *Fgf8* transcription down to zero in a concentration-dependent manner.

*Fgf8* mRNA transcripts have a long half-life of around 2 hours, persisting in cells long after transcription has stopped [27, 32]. This long decay results in a graded distribution of transcript in the spatial maturation domain, with cells proximal to the NMP zone retaining more transcripts that more distal cells. This 2 hour half-life sets the rate constant of degradation to around 0.006 min^−1^ (ln2/2h = 0.693/120min). As the average speed of axis elongation is 3μm/min (from somite 5 to 11; [28]), the decay constant in the spatial maturation domain is 0.002 μm^−1^.

FGF8 protein synthesis is dependent on the concentration of the *Fgf8* transcript within each cell. As FGF8 is synthesized, it diffuses from producing cells at a rate that has been determined experimentally to be around 2 μm^2^/sec [33]. Due to this diffusion, the domain of FGF protein signaling expands beyond the domain of *Fgf8* transcription.

Constant input: *F*_0_(*x*)

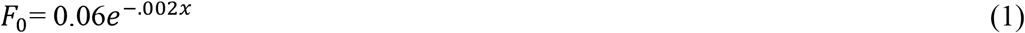

*Fgf8* mRNA transcription: *F*_*m*_(*t*)

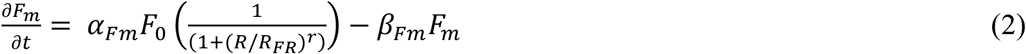

FGF8 translation: *F*(*x*, *t*)

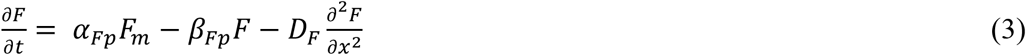

, where,

*Fgf8* mRNA transcription rate constant [27, 29] *α*_*Fm*_ = 1/min

*Fgf8* mRNA half-life [27] *β*_*Fm*_ = 0.006/min.

FGF8 translation rate constant [27] *α*_*Fp*_ = 0.3/min

FGF8 degradation rate constant [27] *β*_*Fp*_ = 0.005/min

FGF8 diffusion constant [33] *D*_*F*_ = 120 μm^2^/min

Hill constant, *Fgf8* inhibition by RA *R*_*RF*_ (see **Table 1**)

#### WNT8C PRODUCTION

*Wnt8c* transcription is stimulated by FGF pathway activity and is indirectly blocked by RA inhibiting *Fgf8* transcription [23]. In chick embryos, *Wnt8c* expression domain significantly overlaps and extends far beyond *Fgf8* mRNA domain up to the last formed somite (**Fig 1A**; [23, 34]), suggesting that very low levels of FGF8 can activate *Wnt8c* [23, 34]. By contrast, in the NMP zone, *Wnt8c* mRNA level are one-third of the *Fgf8* mRNA level [35], which suggest that transcription rate constant of *Wnt8c* are low and saturate quickly [35]. Once synthesized, WNT8C diffuse from its site of synthesis throughout the spatial maturation domain at a low diffusion rate [36].

*Wnt8c* mRNA transcription: *W*_*m*_(*t*)

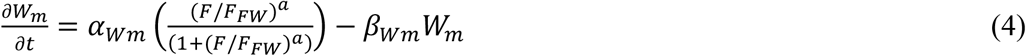

WNT8C translation: *W*(*x*, *t*)

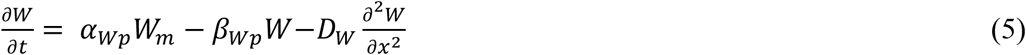

, where,

*Wnt8c* mRNA transcription rate constant [23, 34] *α*_*Wm*_ = 0.1/min

*Wnt8c* mRNA half-life constant [23, 34] *β*_*Wm*_ = 0.03/min

WNT8C translation rates constant [27] *α*_*Wp*_ = 0.3/min

WNT8C degradation rates constant[27] *β*_*Wp*_ = 0.01/min

WNT8C diffusion rate rates [36] *D*_*W*_ = 10 μm^2^/min

Hill constant, *Wnt8c* activation by FGF8 *F*_*FW*_ (see **Table 1**)

#### RA PRODUCTION

RA is synthesized in somites by the enzyme RALDH2 [37]. *Raldh2* transcription is restricted to somites as this is the only region where activation by the WNT8C pathway can overcome FGF8-dependent repression [23]. Parameters for RALDH2 production and degradation were equivalent to those in other models [38]. We assumed that once RALDH2 is produced, RA synthesis initiates without delay. Once produced, RA diffuses into undifferentiated neural and mesodermal tissues at an estimated rate of 18 μm^2^/sec or about 1080 μm^2^/min [38]. At the caudal end of the embryo, RA is degraded by the enzyme CYP26A, whose transcription is under FGF8 regulation [21].

*Raldh2* mRNA transcription: *R*_*m*_(*t*)

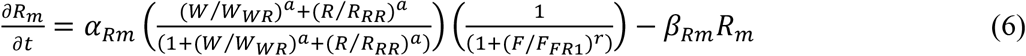

RA production (as modeled by RALDH2 translation): *R*(*x*, *t*)

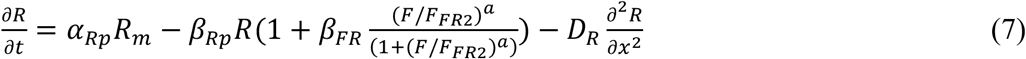

, where,

*Raldh2* mRNA transcription rate constant [27] *α*_*Rm*_ = 1/min

*Raldh2* mRNA half-life constant [27] *β*_*Rm*_ = 0.03/min

RALDH2 translation rates constant [27] *α*_*Rp*_ = 0.3/min

RALDH2 degradation rates constant [27] *β*_*Rp*_ = 0.025/min

RA estimated diffusion rate *D*_*R*_ = 1200 μm^2^/min

FGF dependent RALDH2 degradation constant *β*_*FR*_ = 6/min

Hill constant, *Raldh2* induction by WNT8C *W*_*WR*_ (see **Table 1**)

Hill constant, *Raldh2* induction by RA (autoregulation) *R*_*RR*_ (see **Table 1**)

Hill constant, *Raldh2* repression by FGF8 *F*_*FR*1_ (see **Table 1**)

Hill constant, RA degradation by FGF8-induced CYP26A *F*_*FR*2_ (see **Table 1**)

### Equations modeling the transcript factors interactions network

We used differential equations to simulate the transcription factor network (**Fig 1C**), using as inputs the FGF8, WNT8C and RA output levels obtained in the signaling simulation (represented in the equations with the letters F, W and R, respectively). Network interactions are described in the result section and are supported by previous publications (reviewed in [12]).

*T/Bra* mRNA: *T*_*m*_(*t*)

(Synthesis activated by FGF8 and inhibited by SOX2.)

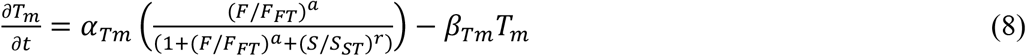

T/BRA protein: *T*(*t*)

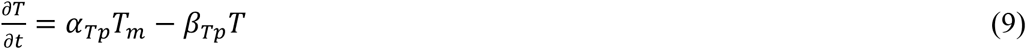

*Sox2* mRNA: *S*_*m*_(*t*)

(Synthesis activated by FGF8 and RA and inhibited by T/BRA.)

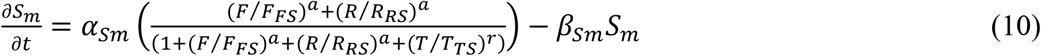

SOX2 protein: *S*(*t*)

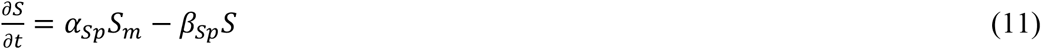

*Nkx1.2* mRNA: *NK*_*m*_(*t*)

(*Nkx1.2* is activated by WNT8C and inhibited by a CDX4-dependent factor X and NKX1.2.)

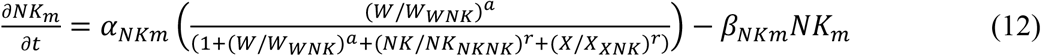

NKX1.2 protein: *NK*(*t*)

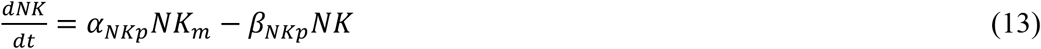

*Cdx4* mRNA: *C*_*m*_(*t*)

(*Cdx4* is induced by FGF8 and WNT8C and inhibited by a PAX6-dependent factor Y.)

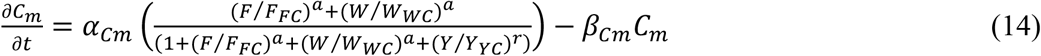

CDX4 protein: *C*(*t*)

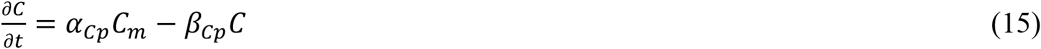

Factor *X* mRNA: *X*_*m*_(*t*)

(Factor *X* is induced CDX4. We have assumed that *X* is inhibited by high FGF8 levels since repression of *Nkx1.2* by CDX4-dependent Factor X is not effective in NMP zone.)

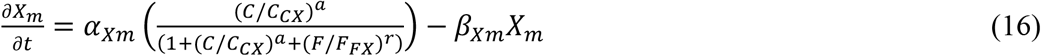

X protein: *X*(*t*)

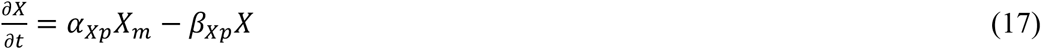

*Pax6* mRNA: *P*_*m*_(*t*)

(*Pax6* is induced by CDX4 and RA working cooperatively, and is inhibited by NKX1.2)

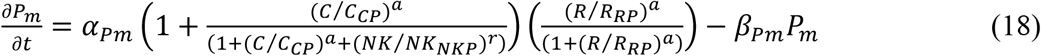

PAX6 protein: *P*(*t*)

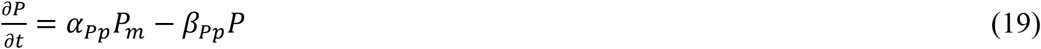

Factor *Y* mRNA: *Y*_*m*_(*t*)

(Synthesis activated by PAX6, and inhibited by FGF8.)

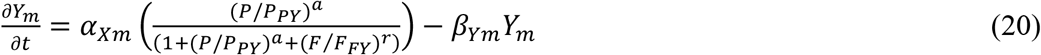

Y protein: *Y*(*t*)

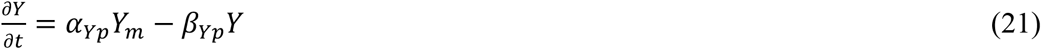

*Ngn2* mRNA: *N*_*m*_(*t*)

(Synthesis activated by PAX6 and inhibited by factor X.)

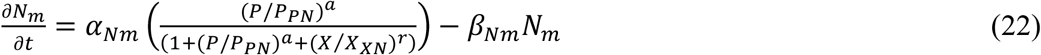

Name definition and values for the Hill constants used in the transcription factor network are found in **Table 2**. For all these transcription factors, the rate constants of mRNA and protein synthesis and degradation have not been determined experimentally. Hence, all the values are kept similar based on values used in published models [27, 29]. The only exception was CDX4, as CDX proteins are known to have increased stability [39].

Constant for mRNA synthesis/degradation: *α*_*im*_ = 1/ min *β*_*im*_ = 0.03/ min

Constant for protein synthesis/degradation: *α*_*ip*_ = 1/ min *β*_*ip*_ = 0.2/ min CDX4

CDX4 constant for protein synthesis/degradation *α*_*cp*_ = 1/ min *β*_*cp*_= 0.05/ min

**Table 2:**
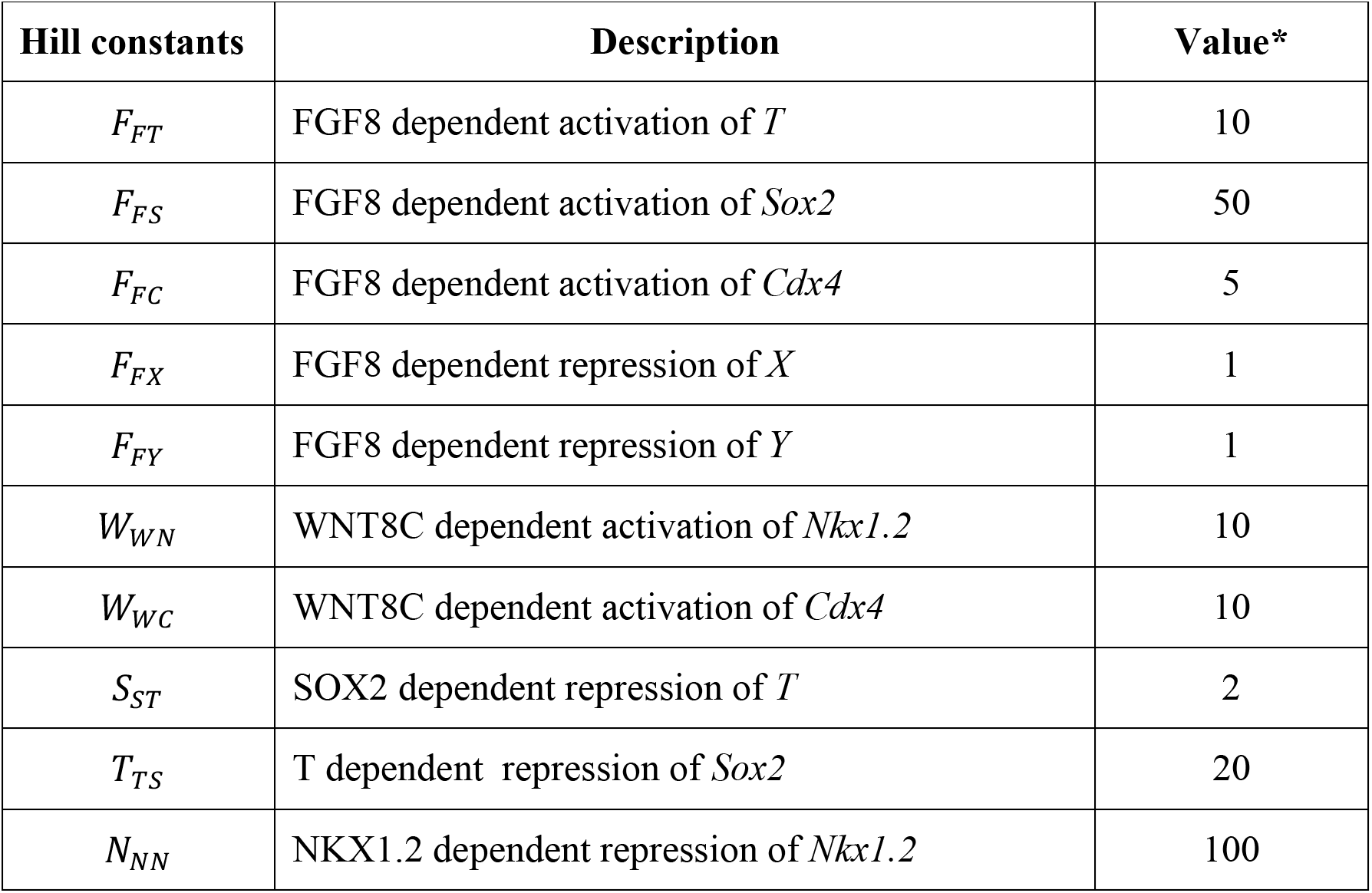
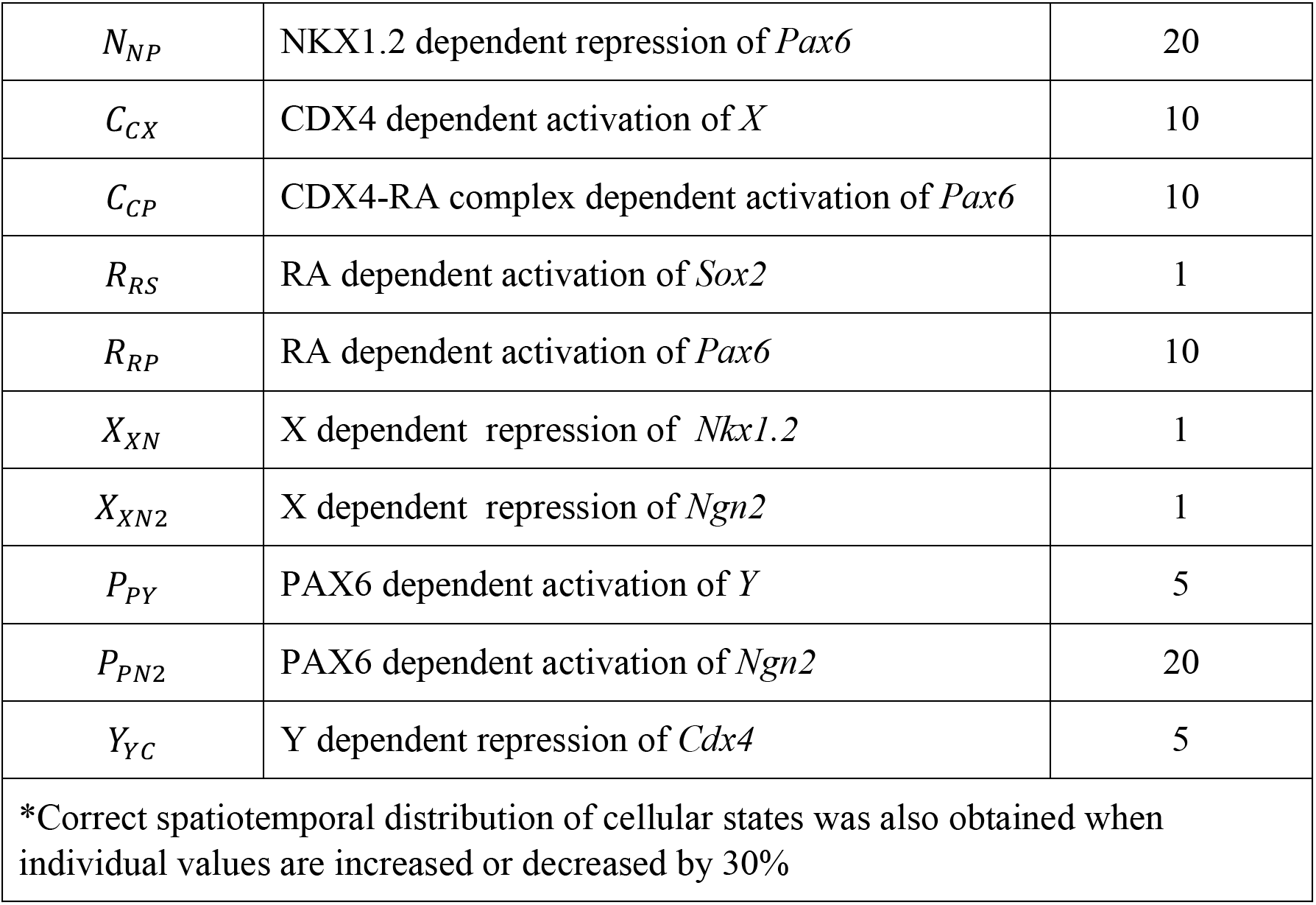
Hill constant for correct spatiotemporal distribution of cellular states.

|  |  |  |
| --- | --- | --- |
| **Hill constants** | **Description** | **Value\*** |
| $F_{FT}$ | FGF8 dependent activation of $T$ | 10 |
| $F_{FS}$ | FGF8 dependent activation of $Sox2$ | 50 |
| $F_{FC}$ | FGF8 dependent activation of $Cdx4$ | 5 |
| $F_{FX}$ | FGF8 dependent repression of $X$ | 1 |
| $F_{FY}$ | FGF8 dependent repression of $Y$ | 1 |
| $W_{WN}$ | WNT8C dependent activation of $Nkx1.2$ | 10 |
| $W_{WC}$ | WNT8C dependent activation of $Cdx4$ | 10 |
| $S_{ST}$ | SOX2 dependent repression of $T$ | 2 |
| $T_{TS}$ | T dependent repression of $Sox2$ | 20 |
| $N_{NN}$ | NKX1.2 dependent repression of $Nkx1.2$ | 100 |
| $N_{NP}$ | NKX1.2 dependent repression of *Pax6* | 20 |
| $C_{CX}$ | CDX4 dependent activation of *X* | 10 |
| $C_{CP}$ | CDX4-RA complex dependent activation of *Pax6* | 10 |
| $R_{RS}$ | RA dependent activation of *Sox2* | 1 |
| $R_{RP}$ | RA dependent activation of *Pax6* | 10 |
| $X_{XN}$ | X dependent repression of *Nkx1.2* | 1 |
| $X_{XN2}$ | X dependent repression of *Ngn2* | 1 |
| $P_{PY}$ | PAX6 dependent activation of *Y* | 5 |
| $P_{PN2}$ | PAX6 dependent activation of *Ngn2* | 20 |
| $Y_{YC}$ | Y dependent repression of *Cdx4* | 5 |
| *Correct spatiotemporal distribution of cellular states was also obtained when individual values are increased or decreased by 30% |  |  |

## Results

### FGF-WNT-RA signaling interaction network can drive signaling switch

In order to model the transcription factor network responsible for spinal cord cell maturation (**Fig 1C**), we first simulated the signaling dynamics between FGF, WNT and RA driving the system [13, 23]. Although several partially redundant FGF and WNT factors are transcribed within and around the caudal neural plate [23, 40], for simplicity, we focused on the factors shown to have the most influence on the system, FGF8 and WNT8C [23]. *Fgf8* is transcribed in the caudal stem zone (**Fig 1A**), where it activates *Wnt8c* transcription [23] and represses RA activity via two mechanisms: by inhibiting transcription of the RA synthesis enzyme *Raldh2* and activating transcription of the RA degradation enzyme *Cyp26a* [21]. FGF8 inhibition of RA production is circumvented rostrally by *Fgf8* mRNA decay [32] and by WNT8C, which stimulates RA production by outcompeting FGF8-mediated *Raldh2* repression [23]. Once *Raldh2* induction has occurred in nascent somites, its expression is maintained through unknown mechanisms even in the absence of WNT activity [23]. For simplification, our model assumes that RA maintains *Raldh2* transcription through positive autoregulation [41]. RA produced by somites then diffuses caudally and inhibit *Fgf8* transcription [21, 42]. These interactions give rise to an extended negative feedback loop between FGF and RA (**Fig 2A**).

Cell proliferation in the stem zone extend the vertebrate body axis caudally by producing the cells that, upon maturation, give rise to the trunk and tail [8]. To simulate the tissue’s caudal ward movement, the signaling interactions were confined to a caudally moving spatial maturation domain of constant length extending rostrally from the stem cell zone to the anterior boundary of the most recently formed somite (**Fig 2B**). Thus, from the perspective of the caudal end, the moving spatial maturation domains appears stationary. To simulate the interactions between signaling factors, we used partial differential equations that integrated synthesis, degradation, and diffusion constant through interaction parameters or Hill constants. The Hill constant of a given reaction is the concentration of a factor involved in transcription or signaling at which the rate of reaction regulated by the factor is half of the maximum possible rate. Hence, Hill constant is inversely related to the affinity of a factor for its target and can act as a measure of factor’s interaction strength (**Fig S1**).

To understand the possible behaviors that could originate from the extended FGF-WNT-RA network, we analyzed the system’s output after systematically changing the signaling inputs and the strength of interaction between components (strong Hill constant =0.1 to weak Hill constant =100). By varying the interaction strength between FGF, WNT and RA components we obtained various temporal signaling information profiles that we grouped into four broad behaviors: FGF-dominance, FGF-RA balance, FGF-RA switch, and RA aberrant/oscillatory.

#### FGF8 dominance

In a system where FGF8 repression of *Raldh2* transcription outweighs RA repression of *Fgf8* transcription, the interactions do not result in appreciable RA production (e. g., **Table 1-I; Fig 2C, Fig S2A**). Such a system would lead to maintenance of pluripotent stem progenitor cells without differentiation.

#### FGF8-RA balance

FGF8, WNT8C and RA signaling domains balance each other and settle on a stable steady state profile (**Table 1-II; Fig 2D, Fig S2A, B**). Such steady state is achieved when the activating and repressive interactions of the system reach an equilibrium. In these conditions, the regions of FGF8 and RA activities are restricted to domains that maintain the same distance from one.

#### FGF-RA switch

The most interesting behavior obtained from simulation is where the system starts with an *Fgf8* mRNA gradient and ends with RA activity gradient over the entire spatial domain (**Table 1-III; Fig 2E, Fig S2A**). This behavior simulates a system that starts with a caudally located stem cell zone and a field of undifferentiated cells that is gradually converted, in a rostral to caudal direction, to a field of differentiate cells. Significantly, this differentiation process is the mechanism by which axial elongation is thought to cease in embryos [31, 43]. The rate at which the FGF8-to-RA transition occurs, and hence differentiation, is modulated by the strength of mutually repressive FGF-RA interactions (**Table 1-IV through VI**; **Fig 3**).

**Fig 3.**
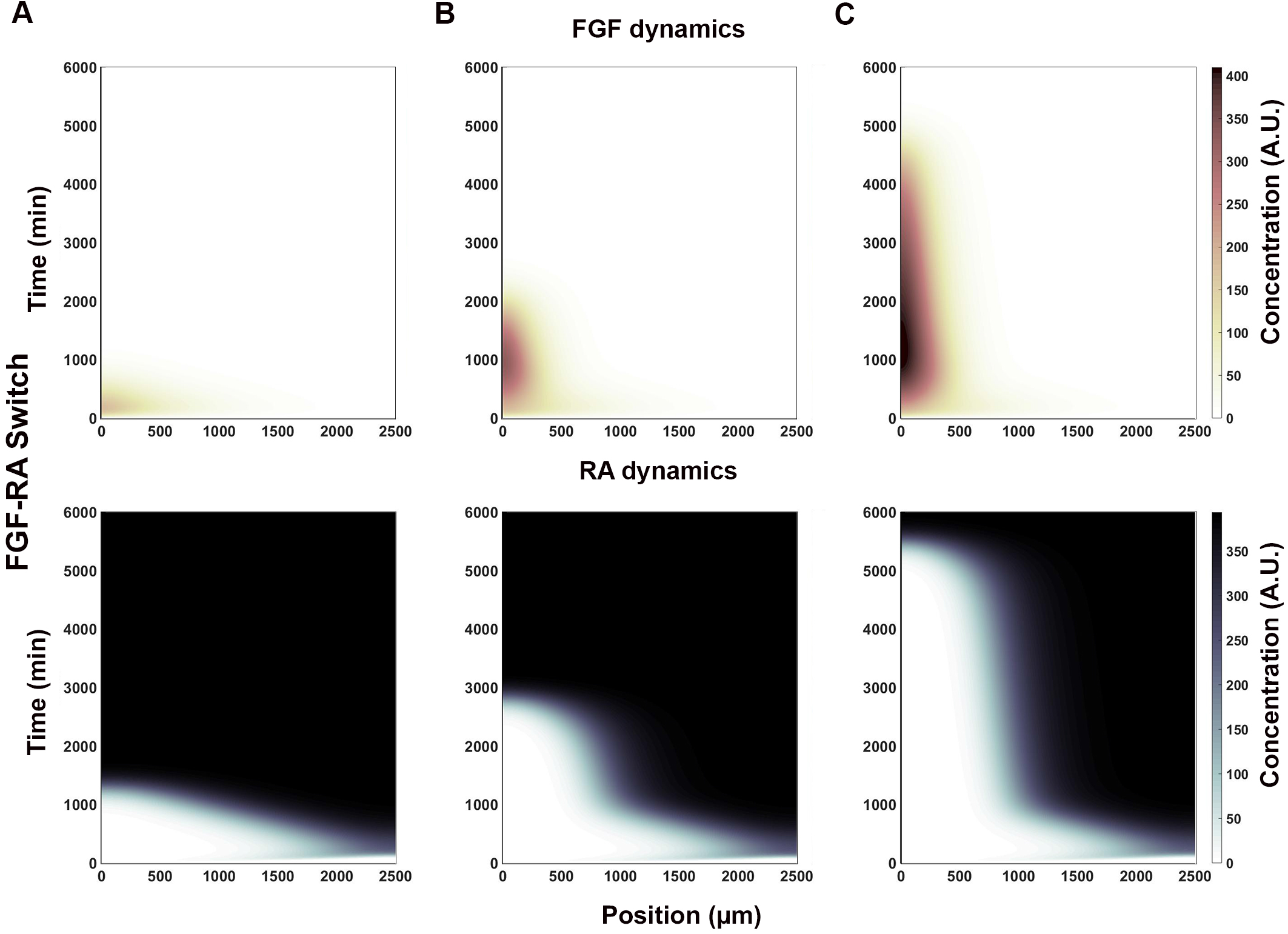
RA inputs strength determine FGF-RA switch rate of conversion. The strength by which WNT8C stimulates (W_WR_) and FGF8 represses (R_RF_) *Raldh2* transcription determines RA’s spatial profile over time (x and y axes, respectively). (**A**) Fast FGF-RA switch (t<1500 min) results from relatively moderate activation and very strong repression inputs (W_WR_=0.5, R_RF_=0.1). (**B**) Intermediate FGF-RA switch (t<3000 min) results from relatively strong activation and moderate repression inputs (W_WR_=0.2, R_RF_=0.3). (**C**) Slow FGF-RA switch (t<5500 min) results from moderate activation and repression inputs (W_WR_=0.5, R_RF_=0.45). FGF and RA heat map scale is shown on the right (arbitrary units; A. U.).

#### RA aberrant/oscillatory

Some parameters in the FGF-WNT-RA interaction system lead to an oscillation in RA levels that did not match the behavior of the system *in vivo*. These oscillations occurred when Hill constants for RA inputs were weak, particularly for the RA-dependent autoregulation of *Raldh2* production (**Table 1-VII, VIII**; **Fig S3**). In some cases, the system produced a discrete burst of RA at the position where the FGF-RA switch was observed, to then return to produce FGF (**Table 1-VII**; **Fig S3A**). In other cases, the burst of RA separated the caudal area of FGF production from a rostral area where FGF and RA production alternated in an oscillatory manner (**Table 1-VIII**; **Fig S3B**).

Altogether, our results show that the FGF8-WNT8C-RA interaction network postulated by Olivera-Martinez and colleagues [23] can indeed give rise to a signaling switch that travels caudally during the elongation of the embryonic axis. The behavior of the switch depends on several interaction parameters that, in coordination, regulate the position and size of the region of cell differentiation.

### FGF-WNT-RA signaling switch and transcription factor network establish areas of pluripotency, early and late differentiation

To simulate the dynamics of the transcription factor network, we integrated the transcription (**Fig 1C**) and signaling (**Fig 2A**) networks into a single supra-network (**Fig 4A**). We then used the FGF-RA balance profile output as the input for the system (**Fig 2D, 4B**), as it most closely resembles the distribution of signaling activity during the steady state period of embryo growth (**Fig 1A**). In this simulation, we followed the transcriptional profile of cells as they are born caudally at t=0 and at subsequent times are displaced rostrally by the appearance of new cells. During their rostral displacement, cells move away from the stem cell zone and the source of FGF and WNT production (**Fig 4C**). As FGF/WNT level decrease, RA levels increase following the FGF-RA balance profile simulation (**Fig 2D, 4B**). These changes in spatial signal information are the drivers for transcription factor expression. Since the cells are arranged spatially from caudal to rostral in order of birth, the temporal changes in transcription factors give rise to spatial changes in profiles.

**Fig 4.**
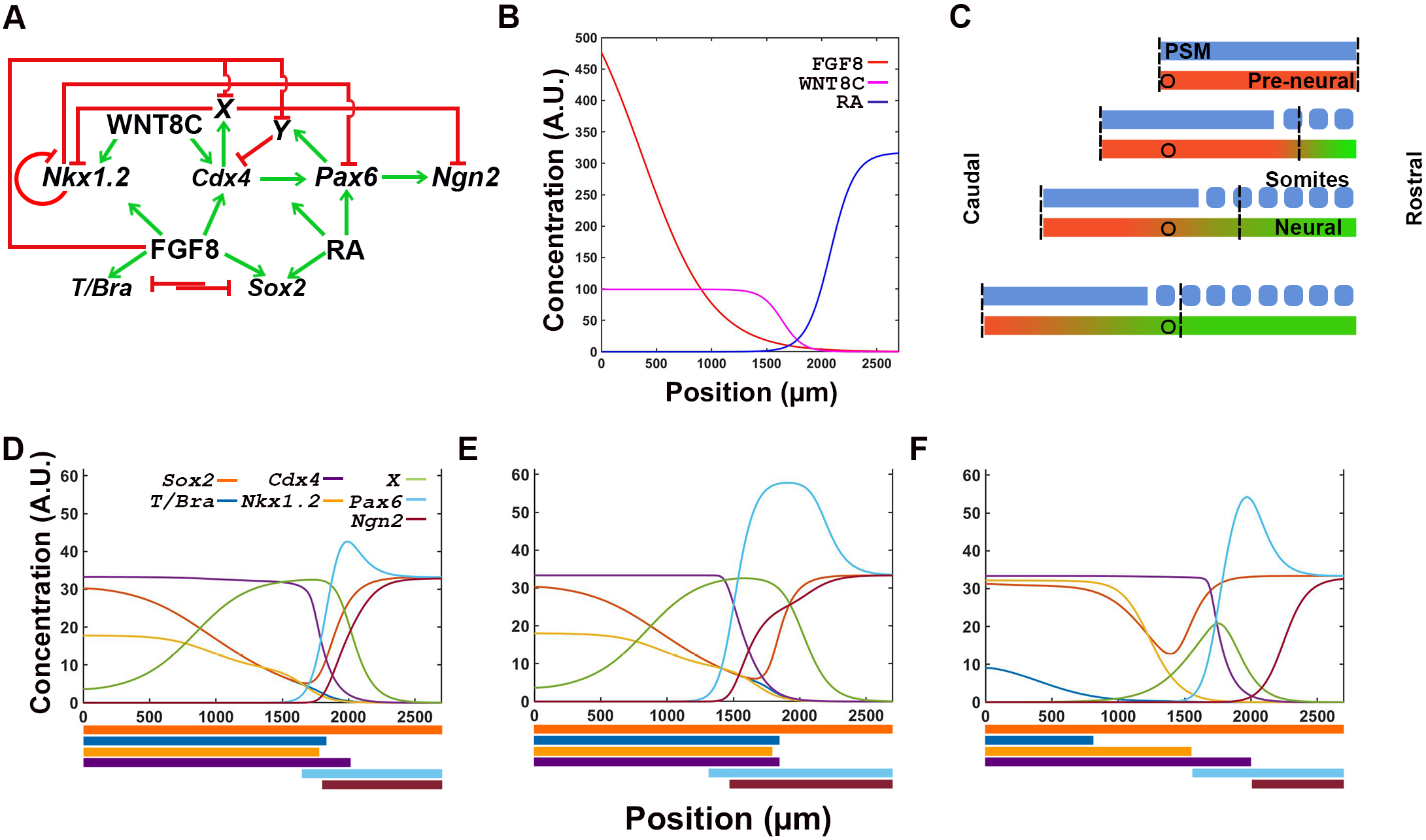
Strength of interactions between transcription factors determine the response of the network to signal information. (**A**) Integrated transcription and signaling interaction network, including hypothetical factors X and Y, predicted from experimental data from reference [12]. (**B**) Reference FGF-RA switch input used for simulations (from **Fig 2D**). (**C**) Location of reference cell (circle) within the spatial maturation domain of signaling activity (vertical dashed lines). Cell remains in the location where it was born as the spatial maturation domain is displaced caudally (as in **Fig 2B**). (**D-F**) Transcription profile of cell with all interactions equally moderate (D; Hill constants=20), equally strong (E; Hill constants=2), and variable (F; as defined in **Table 2**). Output of simulation in F most closely resembles the staggered distribution of transcription factors observed in embryos (**Fig 1B**; [12]). Colored bars at the bottom of each graph represent the spatial domain of gene transcription: *Sox2* in dark orange; *T/Bra* in dark blue; *Nkx1.2* in light orange; *Cdx4* in purple; *Pax6* in light blue; and *Ngn2* in maroon.

The transcription output of the system depends on signaling inputs and transcription factor interactions. Signals regulate the transcription factor network at two distinct key points. The first point of regulation is towards the caudal end of the embryo, where FGF8 and WNT8C, alone or in combination, are required for *T (Bra), Sox2, Nkx1.2* and *Cdx4* transcription [14, 44–48]. Independently of signaling inputs, *Nkx1.2* transcription is negatively regulated by CDX4 and its own protein product [12, 46]. The second point of regulation is towards the rostral end of the neural plate, where RA cooperates with CDX4 to activate the early differentiation gene *Pax6* [12, 49]. *Pax6* is also negatively regulated by NKX1.2 [12, 50]. In contrast, transcription of late differentiation gene *Ngn2* is activated by PAX6 and repressed by CDX4 [12, 22]. Given that CDX4 is an activator and not a repressor [51], our model invokes two putative repressors *X* and *Y* to mediate the repressive activity of CDX4 on *Nkx1.2* and *Ngn2* [12]. These hypothetical repressors are assumed to be inhibited by FGF8 [12].

Signal and transcription factor network were able to generate correct gene transcription profile in many but not all instances (**Fig 4D-F**), indicating that only under certain parameter restrictions could the network recapitulates embryonic events. In principle, the spatial dynamics of the signal interactions network should be sufficient to activate transcription factor network components in the correct spatiotemporal sequence: high FGF caudally would promote pluripotency while high RA rostrally would promote differentiation, with cross-repressive interaction between pathways maintaining the domains separate at opposite ends of the tissue. However, if all the interactions in the network are equally moderate (Hill constants =20, **Fig 4D**) or equally strong (Hill constants =2, **Fig 4E**), then the network does not result in the proper spatial resolution of temporal states. In both cases, transcription of the mesoderm marker *T/Bra* is not restricted to the caudal end, but instead, it is detected throughout the caudal two thirds of the tissue, partially overlapping with *Pax6* and *Ngn2* gene transcripts (**Fig 4D, E**). Only a subset of interaction strengths give rise to correct spatial order of identities (**Table 2; Fig 4F**). The values of the interactions strengths that generate proper spatial distribution of transcripts could be increased or decreased by 30%. These values define a parametric space where the model is operational and highlights its robustness **(Fig S4).** These results suggest that signaling inputs encodes the information required for specifying different cell maturation states, but that it is the transcription factor network what determines the spatial distribution and organization of maturation states cell along the caudal-to-rostral length of the tissue.

### The transcription network executes the spatiotemporal information provided by the signaling factor network

To further evaluate the contribution of signaling and transcription factors networks on cell maturation events, we tested the effect of disrupting individual network nodes on transcription readouts. First, we tested the response of the transcription network to signaling noise. In simulations, both periodic disturbance (**Fig 5A**) and random noise (**Fig 5B**) were well tolerated by the transcription network without any distortions in the spatiotemporal resolution of the cellular states. Unexpectedly, introduction of random noise resulted in better separation of maturation (Cdx4^+^, Pax6^+^, Ngn2^−^) and late differentiation (Cdx4^−^, Pax6^+^, Ngn2^+^) states (**Fig. 4F, 5B**). This suggest that transcription network has built in robustness to extrinsic disturbances.

**Fig 5.**
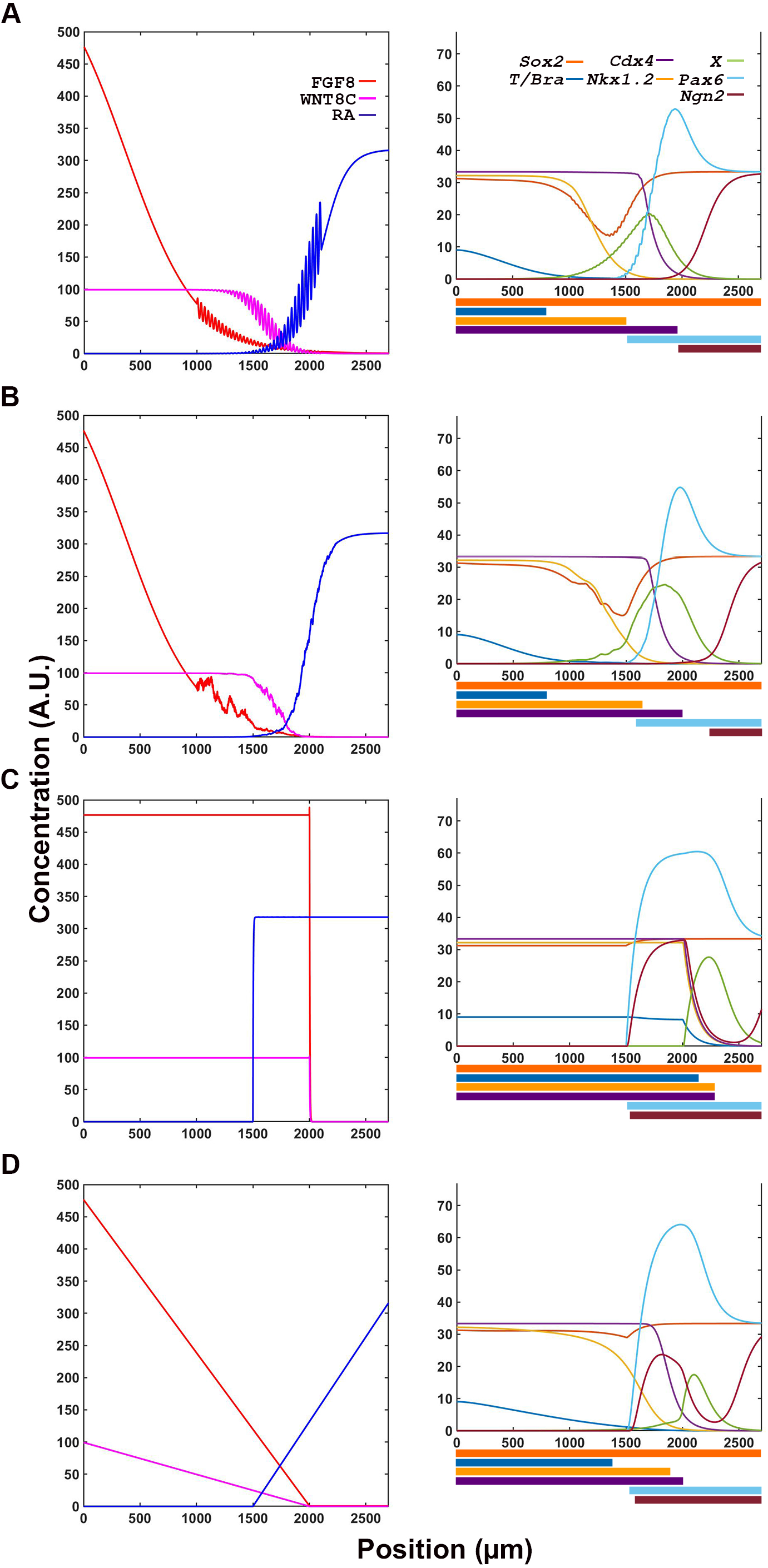
Transcription factor network is resilient to small and moderate alterations in signaling information. (**A, B**) Transcription factor output is not affected by oscillatory (A) or random noise (B) in signaling inputs, as outputs are comparable to those obtained in conditions without noise (Fig 4F). (**C, D**) Large changes in signaling input such as discreet Boolean (C) and linear gradient (D) changes transcription factor expression domains.

Next, we evaluated the role of signaling gradients in determining the spatiotemporal resolution of downstream targets’ transcriptional domains. Replacing the exponential gradient of the signaling factors with a Boolean switch (**Fig 5C**) or a linear gradients (**Fig 5D**), resulted in loss of proper resolution of transition zones. Thus, changes in the spatial information contained in the signaling network changes the transcription network readouts. This confirms that the spatial information is encoded in the signaling and not in transcription factor network.

We previously proposed a central role of CDX4 in regulating maturation of NPs in the pre-neural tube. To theoretically test CDX4 role in transcription network regulation, we removed, increased or introduced noise to *Cdx4* transcription and evaluated the network’s transcription profile output (**Fig 6**). When *Cdx4* was removed from the simulation, *Nkx1.2* transcription expanded rostrally, overlapping significantly with the expression of differentiation markers *Pax6* and *Ngn2* (**Fig 6A**). This phenomenon is opposite to what is observed experimentally, were downregulation of CDX4 activity using an ENRCDX4 repression construct results in downregulation of *Nkx1.2* [12] (discussed below). Conversely, when the levels of *Cdx4* were increased in the simulation, *Nkx1.2* expression domain shifted caudally, away from *Pax6* expression domain, and rostral cells did not activate the late differentiation gene *Ngn2* (**Fig 6B**), in agreement with experimental results [12]. Thus, removing or increasing *Cdx4* transcription affects the spatial relationship between early specification gene *Nkx1.2* and neural differentiation gene *Ngn2*, indicating that CDX4 function in the network is key to establish a transition zone between pluripotency and differentiation states. This function of CDX4 is robust, as introduction of transcriptional noise still produce the expected gene expression profile with only minute changes in the position of boundary transitions (<+/−30μm; **Fig 6C)**. Thus, with the one exception of Cdx4 removal discussed below, our simulation generally agrees with our *in vivo* observed role of CDX4 in driving NP maturation during early spinal cord development.

**Fig 6.**
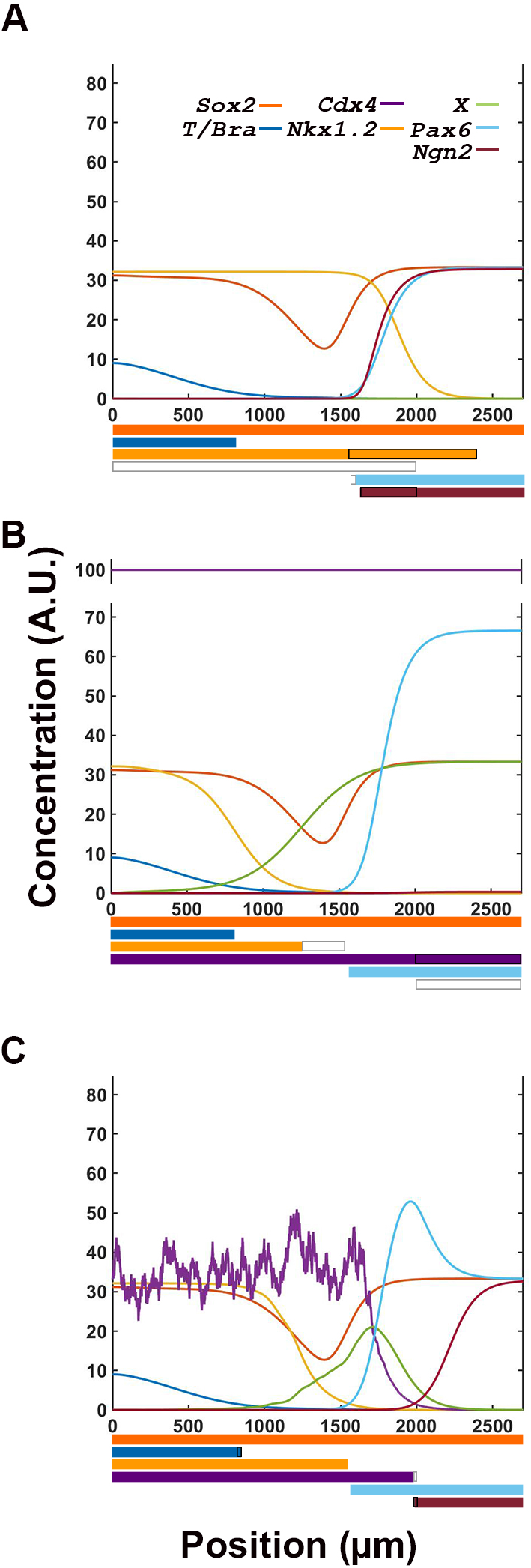
CDX4 is necessary for proper interpretation of signaling inputs by the transcription factor network. (**A**) Compared to control simulation (Fig 4F), loss of *Cdx4* expression causes a large rostral expansion of *Nkx1.2* domain, a small reduction in *Pax6* domain and a large caudal expansion in *Ngn2* domain. These changes results in the overlap of stem and differentiation gene expression domains. (**B**) Overexpression of *Cdx4* reduces *Nkx1.2* and eliminates *Ngn2* expression domains, effectively expanding the early and eliminating the late differentiation zones. (**C**) Introduction of random noise in *Cdx4* transcriptional noise has insignificant effects on the system’s spatial expression profiles. In the expression profile bars at the bottom of the graphs, white and black rectangles indicate loss and gain of gene transcription, respectively.

## Discussion

### Signaling factor simulation recapitulates signaling dynamics observed in natural systems

Our simulations describe the possible behaviors the FGF8-WNT8C-RA system can exhibit under various interaction conditions (**Fig 2**). With small variations in interactions’ strength, the system can model behaviors associated with different stages of axial tissue development. In a system where the FGF’s activity dominates over RA’s activity, the simulation most closely resembles the neural tissue at early stages of axial extension, whereas in a system where FGF activity balances that of RA, the simulation resembles the maturation of spinal cord cell that occurs during axial elongation. In contrast, a switch in the system from FGF to RA most closely resembles the processes occurring during termination of body axis extension [9, 28]. Significantly, when the interactions between FGF and RA components are weak, the system oscillates, resembling the oscillations observed between FGF/WNT and the Notch signaling pathway during the process of paraxial mesoderm segmentation [52]. Thus, with small modifications in signal components interaction, one can observe large changes in the behavior of the system equivalent to the changes normally observed in the tissues emerging from the tailbud during axial elongation, the paraxial mesoderm and spinal cord.

We propose a model of vertebrate body extension where modulation of interaction strength between different components of the system (e.g., transcriptionally, post-transcriptionally or epigenetically), could regulate the spatiotemporal dynamics involved in vertebrate body extension. In this model, the time at which the system transitions from FGF dominant, to FGF-RA balance, to RA switch respectively determine the time of tissue induction, elongation and termination. For example, a long period in which the FGF8-RA balance system is operational could explain the elongated axis of vertebrates such as snakes; as long as the FGF8-RA balance system remains operational, the caudal progenitor/stem cell pool will continue to generate tissue and extend the axis. In this scenario, the time at which RA takes over the system to initiate progenitor cell differentiation will determine the axial body length, with a fast FGF-RA switch resulting in a shorter body axis and vice versa.

### Transcription network simulations recapitulate the cell state transitions observed in the caudal neural plate

Results from simulations support a role for CDX in coordinating upstream signaling factors with downstream transcription network components involved in spinal cord neural maturation. In the present model, CDX4 functions to separate caudal stem cell populations (*Nkx1.2*^+^ *Pax6-Ngn2*^−^) from rostral differentiating cells (*Nkx1.2*^−^*Pax6*^+^ *Ngn2*^+^) by establishing a transition zone. This is achieved by CDX4 repressing the bipotency gene *Nkx1.2* and the late differentiation gene *Ngn2*, and by activating the early differentiation gene *Pax6*. In simulations, high levels of *Cdx4* transcription resulted in downregulation of CDX4 repressed genes (**Fig 6B**): *Nkx1.2* transcription domain shifted caudally and *Ngn2* transcription was lost. In these conditions, only the caudal expression of *Nkx1.2* was retained due to its high dependence on WNT stimulation [46]. Increasing *Cdx4* transcription did not affect the expression domain of *Pax6,* as transcription of this gene is also dependent on RA secreted from somites [12, 49]. Together, the changes in *Nkx1.2* and *Ngn2* transcription induced by CDX4 overexpression effectively increase the size of the transition zone. The same way that premature activation of differentiation signals has been predicted to cause shortening of the embryonic axis [31], a greater separation of stem cell and differentiation signals is predicted to cause axial lengthening. These predictions would need to be tested experimentally.

Significantly, results obtained by simulating loss of *Cdx4* activity (**Fig 6A**) did not fully match experiments done *in vivo*. With respect to differentiation genes, the network recapitulates the *in vivo* results: *Pax6* transcription was not affected due to dependence of this gene on RA [12, 49], whereas *Ngn2* transcription was upregulated because this gene is normally repressed by CDX4 [12]. In contrast, with respect to the NMP marker *Nkx1.*2, loss of *Cdx4* caused an anterior expansion of *Nkx1.2* expression domain that was not observed experimentally. This discrepancy can be attributed to the use of a dominant negative form of CDX4 instead to knockout allele to downregulate the activity of this gene *in vivo* (ENRCDX4; [12]). Dominant-negative ENRCDX4 works by outcompeting endogenous CDX4 from binding to its target genes and repressing their transcription [53]. This approach is different than not having CDX4 protein altogether (e. g, through deletion of the gene). Given that our model simulates the loss of CDX4 function and not the active repression of its downstream target genes, this providing a possible explanation for the observed discrepancies between experimental systems. It is also possible that our current understanding of the transcription factor network is incomplete. These two possible explanations are not mutually exclusive, and could be resolved with additional experiments. While additional experiments will be required to fully understand CDX4 function in the NMP zone, even with its limitations, the proposed transcription factor network supports a key role for CDX4 in the segregation of cell states in the nascent spinal cord, providing a robust mechanisms for the progressive maturation of cells during axial elongation.

### Future perspectives

Although the integrated signaling and transcription factor network model presented here provides key information on the transition state drivers underlying neuronal cell maturation, it is clear from experimental and modelling data that the model is far from complete. One key component missing from the model is the feedback control that transcription factors have over the signaling network. Chromatin immunoprecipitation studies comparing anterior versus posterior portions of mouse embryos have shown that CDX2 can regulate *Wnt5a*, *Rspo3* (a WNT pathway component), *Fgf4* and *Fgf8*, indicating that CDX2 contributes to signaling factor regulation and axial growth [54]. Similarly, CDX binding sites are present in *Radh2* intronic enhancer are sufficient to drive reporter gene expression in the caudal end of embryos [55]. Currently, however, lack of quantitative data precluded the incorporation of feedback activities into an integrated network model.

Our modeling results also highlights the importance of signaling factor regulation by components external to the signaling pathways. For example, our model shows that maintenance of RA production is critical for the behavior of the system, as weakening of the Hill constant regulating RA-dependent autoregulation of *Raldh2* production causes the system to transition from balanced to oscillatory (**Table 1-VII, VIII**; **Fig S3**). While, for simplification purposes we assumed that Raldh2 maintenance is dependent or RA, it is likely to be dependent on transcription factors, some of which are part of our transcription network (e.g. CDX; [55]). Understanding the effect that transcription factor network components have over the signaling network will be important for understanding the later stages in neural cell maturation and their subsequent differentiation.

## Author contributions

P.J. and I.S. designed the experiments. P.J. derived the mathematical equations, coded the simulation and performed experiments. P.J. and I.S. analyzed the results and wrote the manuscript.

## Acknowledgements

We thank Dr. Donald DeAngelis for guidance and for sharing his mathematical expertise, and members of the Skromne lab for intellectual discussion. We also thank Dr. K. G. Story (U Dundee, UK), Dr. M. Gouldin (Salk Institute, USA). Dr. F. Medeville (CBI, France), Dr. S. Mackem (NCI, USA), Dr. Y. Marikawa (U Hawaii, USA), Dr. A. V. Morales (Cajal Institute, Spain) and Dr. B. Novitch (UCLA, USA) for generously providing plasmids.

## SUPPLEMENTARY INFORMATION

### Supplementary methods

1. General Hill functions
2. In situ hybridization

### Supplementary figures

**Fig S1.**
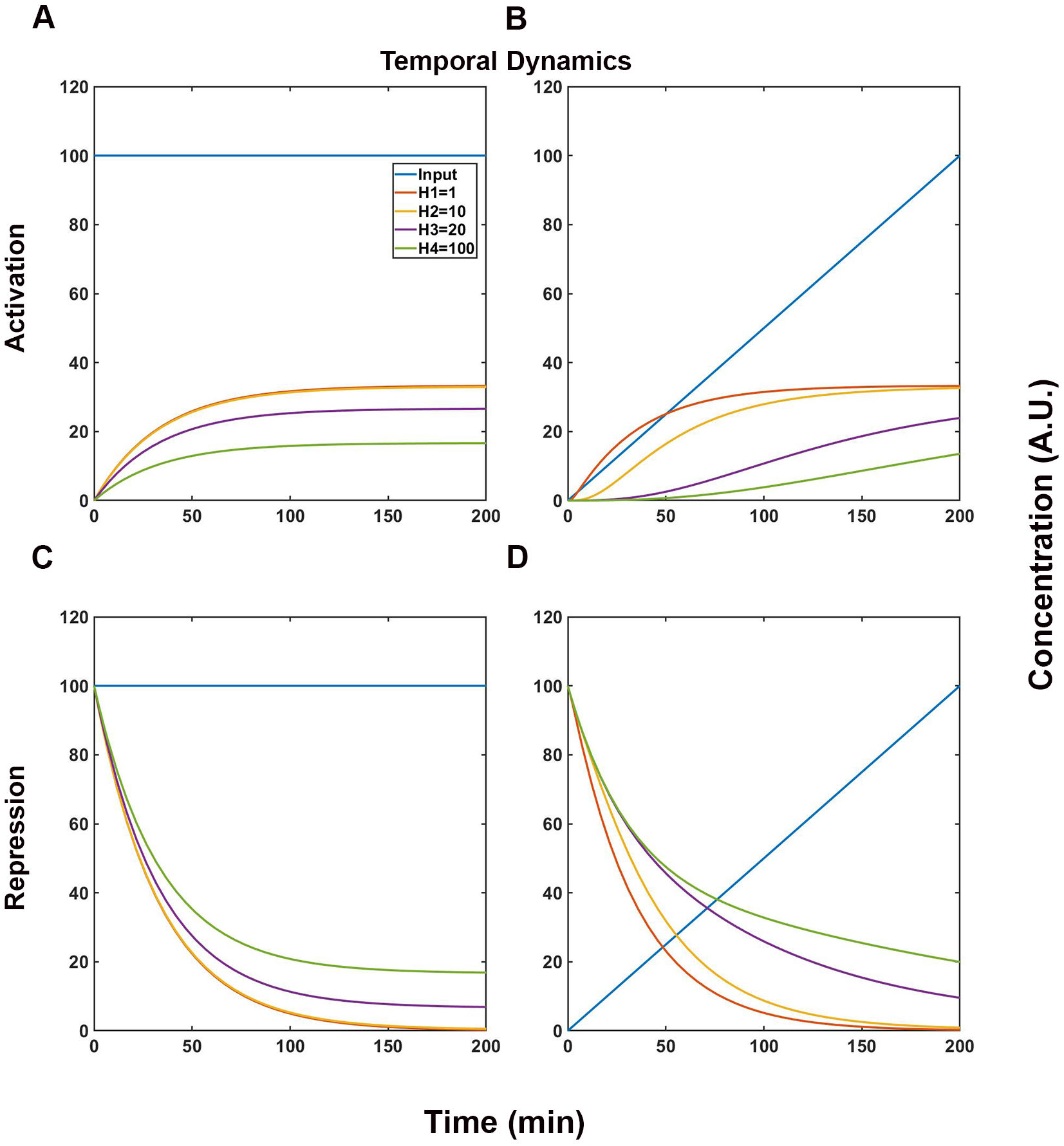
Hill constant determine the strength of response of targets to activators and repressors. Temporal response of targets with different Hill constants to activators and repressors. Inputs are shown in blue and targets with different Hill constants are color coded: H1=1, orange; H2=10, yellow; H3=20, purple; and H4=100, green. **(A-B)** For activators, constant (A) and graded (B) inputs induce targets with smaller Hill constants to higher levels than targets with larger Hill constants. (**C-D**) For repressors, constant (C) and graded (D) inputs reduce targets with smaller Hill constants to lower levels than targets with larger Hill constants. With graded inputs (B, D), larger Hill constants also cause temporal delays in response.

**Fig S2.**
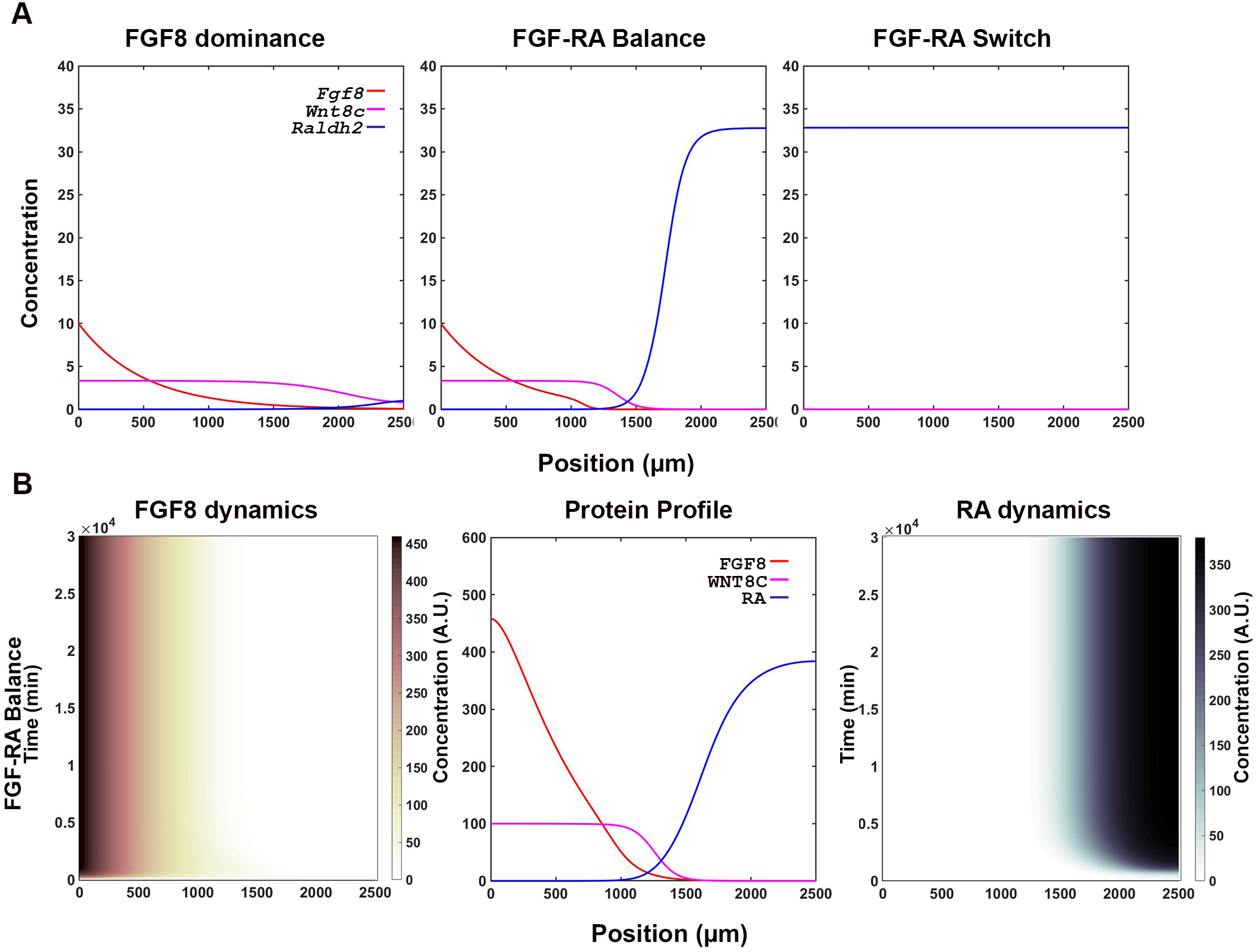
Changes in FGF-WNT-RA signaling interactions results in mRNA profiles parallel protein accumulation and are stable over time. (**A**) Profiles of mRNA transcripts at t=6000 min associated with production of signaling molecules. Transcript and protein profiles are similar (**Fig 2C** left panels). (**B**) Signaling molecule profiles are stable over longer simulation times. An FGF-RA balance simulation that was run for t=30,000 min produced the same profile than a simulation that was run for t=6000 min (**Fig 2C** middle row).

**Fig S3.**
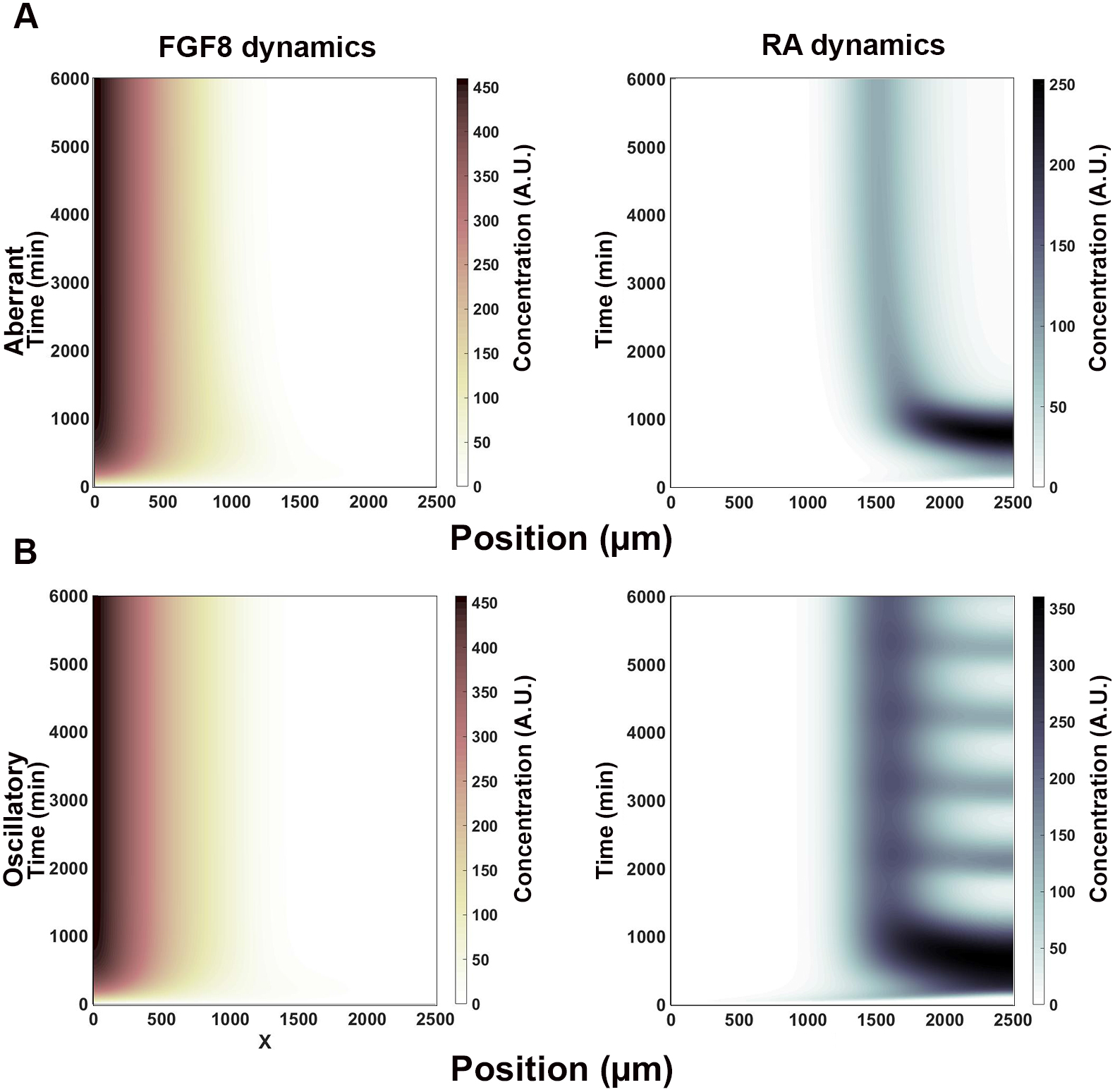
RA positive autoregulation is required for bistability. Reducing RA’s positive effect on *Raldh2* transcription (H= 300 instead of 50) results in aberrant RA, but not FGF, distribution. (**A**) Under these conditions, when FGF affinity to repress *Raldh2* is strong (H=2 instead of 10), a peak of RA production forms at a position in the field where the FGF-RA switch would have occurred (1500-2000 μm). (**B**) When RA repression of *Fgf8* transcription is weakened (H=20 instead of 1), RA production oscillates in the region of cell differentiation (>1500 μm).

**Fig S4.**
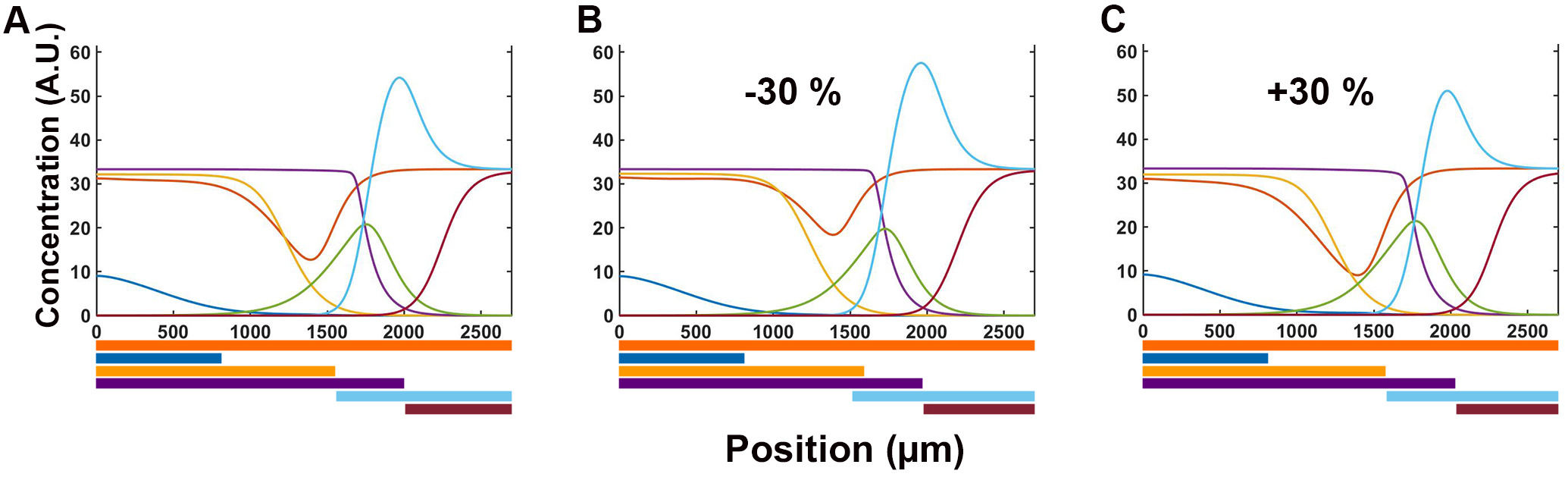
Stability of the transcription profile in the parameter space. Changes in the strength of interactions between transcriptional factors does not drastically affect the transcriptional domain profile. (**A**) Original transcription profile as shown in Fig 4F. (**B, C**) Reducing (B) or increasing (C) all the Hill constants in the interaction network by 30% does not significantly change the spatial profile of gene transcription.

## APPENDIX

**SIGNET.m**: MATLAB code for simulating signaling dynamics.

**TRANSNET.m**: MATLAB code for simulating transcriptional factor dynamics.

## SUPPLEMENTAL INFORMATION

### Supplementary methods

#### Hill functions describing general regulatory interactions

The Hill functions *H*_1_, *H*_2_, *H*_3_, *H*_4_ were used in the transcription and translation equations to describe one of four different types of interactions: (1) inductive, (2) repressive, (3) coordinated and (4) competitive. We generated transcription and translation equations specific for each factor by replacing *H*_1_, *H*_2_, *H*_3_, and *H*_4_ with the functions that represent the regulatory interactions reported experimentally.

##### INDUCTIVE INTERACTIONS

*H*_*i*_, describes the input of one or multiple activators and can take three different forms.

*H*_*i*_ for a single activator is of the form:

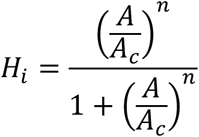

*H*_*i*_ for two activators, when both activators are necessary to stimulate activity (Boolean AND), is of the form:

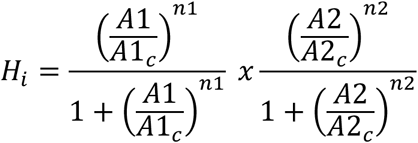

*H*_*i*_ for two activators, when one activator is sufficient to stimulate activity (Boolean OR), is of the form:

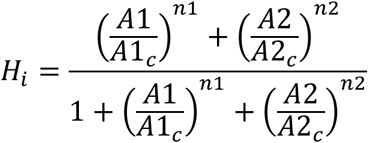

, where, for all these equations, *A*, *A*1, *A*2 are the concentrations of activator proteins; *A*_*c*_, *A*1_*c*_, *A*2_*c*_ are the respective Hill constants; and *n, n1, n2* are the Hill coefficient of cooperativity.

##### REPRESSIVE INTERACTIONS

*H*_*i*_, describes the input of one or multiple repressors and can take three different forms.

*H*_*i*_ for a single repressor is of the form:

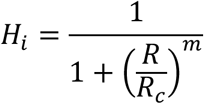

*H*_*i*_ for two repressors, when both repressors are necessary for downregulation (Boolean AND), is of the form:

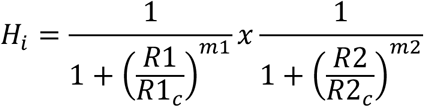

*H*_*i*_ for two repressors, when one repressor is sufficient for downregulation (Boolean OR), is of the form:

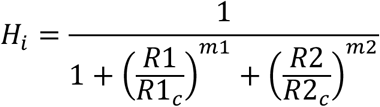

, where, for these equations, *R*, *R*1, *R*2 are the concentration of repressor proteins; *R*_*c*_, *R*1_*c*_, *R*2_*c*_ are the respective Hill constants; and *m, m1, m2* are the Hill coefficients of cooperativity.

##### COORDINATED INTERACTIONS

*H*_*i*_, describes the input of multiple activators and repressors working simultaneously through separate regulatory sites.

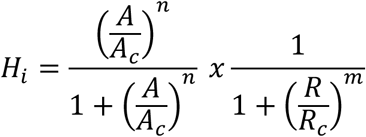

, where, for these equations, *A*, *R* are the concentration of activator and repressor proteins; *A*_*c*_, *R*_*c*_ are the respective Hill constants; and *m, n* are the Hill coefficients of cooperativity.

##### COMPETITIVE INTERACTIONS

*H*_*i*_, describes the input of multiple activators and repressors working simultaneously through the same regulatory sites.

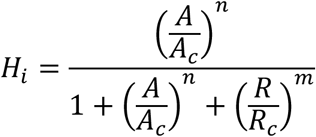

, where, for these equations, *A*, *R* are the concentration of activator and repressor proteins; *A*_*c*_, *R*_*c*_ are the respective Hill constants; and *m, n* are the Hill coefficients of cooperativity.

Hill constants measure the interaction strength between the activators/repressors and their target gene, and are defined as the concentration of the regulator at which the rate of gene activation or repression is half the synthesis rate maximum [1]. The hypothetical range for Hill constants is from zero to infinity, with low Hill values signify strong and fast regulatory activities while high Hill values signifying weak and slow regulatory activities (**Fig S1**).

#### In situ hybridization

Analysis of gene transcription by *in situ* hybridization was done using digoxigenin (DIG)-labeled antisense RNA probes synthesized and hybridized using standard protocol [2], as previously described [3]. Briefly, embryos were harvested at the appropriate stage and fixed with 4% paraformaldehyde diluted in 1x PBS at 4° C overnight. After a series of washes, embryos were exposed overnight in hybridization solution to DIG-labeled antisense RNA probes against *Pax6*, *Nkx1.2*, *Bra*, *Sox2*, *Cdx4 or Ngn2*. mRNA expression was detected using an Alkaline Phosphatase coupled Anti-DIG antibody (Roche) and developing embryos with nitro-blue tetrazolium salt (NBT, Thermo Scientific) and 5-bromo-4-chloro-3-indolyl-phosphate (BCIP, Biosynth) at room temperature until dark purple precipitate deposited revealing the areas of gene transcription. Post-development, embryos were washed with 1x TBST and then fixed in 4% PFA. Processed embryos were photographed using an AxioCam MRc digital color camera mounted on a Zeiss V20 Stereo microscope and processed using Adobe Photoshop (CC2017, Adobe) for size and resolution adjustment and figure preparation.

